# Learning and Imputation for Mass-spec Bias Reduction (LIMBR)

**DOI:** 10.1101/301242

**Authors:** Alexander M Crowell, Casey S Greene, Jennifer J. Loros, Jay C Dunlap

**Affiliations:** Department of Molecular and Systems Biology, Geisel School of Medicine at Dartmouth, Hanover NH, 03755, USA; Department of Systems Pharmacology and Translational Therapeutics, Perelman School of Medicine, University of Pennsylvania, Philadelphia PA 19104, USA; Department of Biochemistry and Cell Biology, Geisel School of Medicine at Dartmouth, Hanover NH, 03755, USA

## Abstract

**Motivation:** Decreasing costs are making it feasible to perform time series proteomics and genomics experiments with more replicates and higher resolution than ever before. With more replicates and time points, proteome and genome-wide patterns of expression are more readily discernible. These larger experiments require more batches exacerbating batch effects and increasing the number of bias trends. In the case of proteomics, where methods frequently result in missing data this increasing scale is also decreasing the number of peptides observed in all samples. The sources of batch effects and missing data are incompletely understood necessitating novel techniques.

**Results:** Here we show that by exploiting the structure of time series experiments, it is possible to accurately and reproducibly model and remove batch effects. We implement Learning and Imputation for Mass-spec Bias Reduction (LIMBR) software, which builds on previous block based models of batch effects and includes features specific to time series and circadian studies. To aid in the analysis of time series proteomics experiments, which are often plagued with missing data points, we also integrate an imputation system. By building LIMBR for imputation and time series tailored bias modeling into one straightforward software package, we expect that the quality and ease of large-scale proteomics and genomics time series experiments will be significantly increased.

## 1 Introduction

In recent years, the scope of whole proteome mass spectrometry (MS) experiments have expanded drastically Chick *et al*., 2016; Weekes *et al*., 2014. In principle, increased sample size can make the expression patterns identified by these experiments more robust. However, increasing sample sizes exacerbate many of the technical limitations of MS. The issues of missing data and batch effects that are common to this experimental platform hamper analyses of dozens of samples Karpievitch *et al*., 2012, and this can be particularly problematic when sequential data sets are collected with the goal of discerning trends, for instance in the field of circadian rhythms.

The first challenge presented by larger scale experiments is the failure to observe peptides in all samples. The abundance of a peptide can go unmeasured in a sample because of low abundance, random sampling or as a function of its sequence Tabb *et al*., 2003; Mallick *et al*., 2006. Many circadian analysis methods, including the best Jonckheere-Terpstra-Kendall based methods, do not support missing data, forcing researchers to remove peptides with any unobserved values or use inferior tests of rhythmicity Hughes *et al*., 2010; Hutchison *et al*., 2015; Mauvoisin *et al*., 2014; Robles *et al*., 2014, 2016; Wang *et al*., 2017a. Because the missingness of peptides can depend on factors other than abundance and random sampling (for example sub-cellular localization or degree of structure), the analysis may in addition be biased. Missingness as a function of abundance can also bias analyses because only the most abundant and least dynamically expressed peptides will be present in a complete-case analysis Gelman and Hill, 2007. If we imagine a dynamically expressed peptide with a modest average expression level, such a peptide will have lower lows of expression than either a higher expressing dynamic peptide or a constitutively expressed peptide with the same mean expression. Such peptides are more likely to be dropped from analyses. The prevalence of missing data also leads to the counterintuitive situation that as the number of replicates increases, fewer and fewer peptides are measured in all samples and available for analysis with best-in-class methods. It is therefore critical to recover at least those peptides missing in only a small fraction of observations before analysis.

Batch effects are the second challenge that larger scale experiments exacerbate. A systematic study of variance in iTRAQ experiments showed that measurements of abundance in most proteins can be expected to exhibit 10-20% relative errors, going as high as 40% for low weight proteins with few detected peptides Hultin-Rosenberg *et al*., 2013. Most modern experimental designs include pooled control samples; however, such pooled controls account only for variability introduced by MS runs and not for bias introduced by sample handling, which has been shown to contribute similar amounts of variability Piehowski *et al*., 2013. Even for the best Jonckheere-Terpstra-Kendall based methods, the addition of 10-20% error meaningfully diminishes the ability to classify circadian expression patterns Hutchison *et al*., 2015. Batch effects are a greater problem as the number of samples increases, first because the number of batches must increase accordingly making relative comparisons in the expression of a given peptide more challenging, and second, because more peptides will be influenced by at least one batch effect. It is therefore critical to be able to model and remove bias trends when analyzing large-scale proteomics data.

Here we provide a toolset developed in response to these issues, Learning and Imputation for Mass Spec Bias Reduction (LIMBR). LIMBR employs a K nearest neighbors (KNN) based imputation strategy that is both non-parametric and has been shown to be highly effective in the context of proteomics data Wang *et al*., 2017b. LIMBR’s bias trend modeling procedure is based on SVA, a proven bias modeling algorithm initially devised for microarray data Leek, 2007; Leek and Storey, 2007b; Leek *et al*., 2012 that has been used extensively and successfully in many systems Parsana *et al*., 2017; Benjamin *et al*., 2017; Wang *et al*., 2016; Lopez *et al*., 2014; Tsang *et al*., 2014 and has also spawned many adaptations Parker *et al*., 2014; Chakraborty *et al*., 2013; Karpievitch *et al*., 2009. LIMBR improves SVA with optimizations specific to large-scale MS experimental designs, particularly in its handling of general and circadian time courses. We report extensive tests on both simulated and experimental data, documenting that LIMBR accurately removes bias trends without requiring information about sample preparation and handling.

### Approach

The need for LIMBR software arose out of technical challenges presented by a very large circadian proteomics time course conducted in *N. crassa*. In brief, cultures of the fungus were synchronized to the same circadian time and released into a circadian free-run; samples were then collected at regular intervals and iTRAQ based MS was performed. This dataset has measurements for 48 hours with a two hour resolution. Two genotypes are measured and all assays are performed with three replicates for a total of 144 experimental samples. With pooled controls, 192 total MS samples were assayed.

Because of the number of replicates and total conditions, only 24% of the total assayed peptides were detected in all samples. Approximately one-third of peptides were missing in almost all samples, one third were present in the majority of samples, and the remainder were roughly uniformly distributed across intermediate levels of missingness (Fig. 1 a). For downstream analysis methods that perform complete-case analysis, 76% of detected peptides would have been discarded. An illustration of the decrease in the number of peptides considered in a complete case analysis as the size of the experiment is increased is shown in Fig. 1 b.

**Fig. 1.**
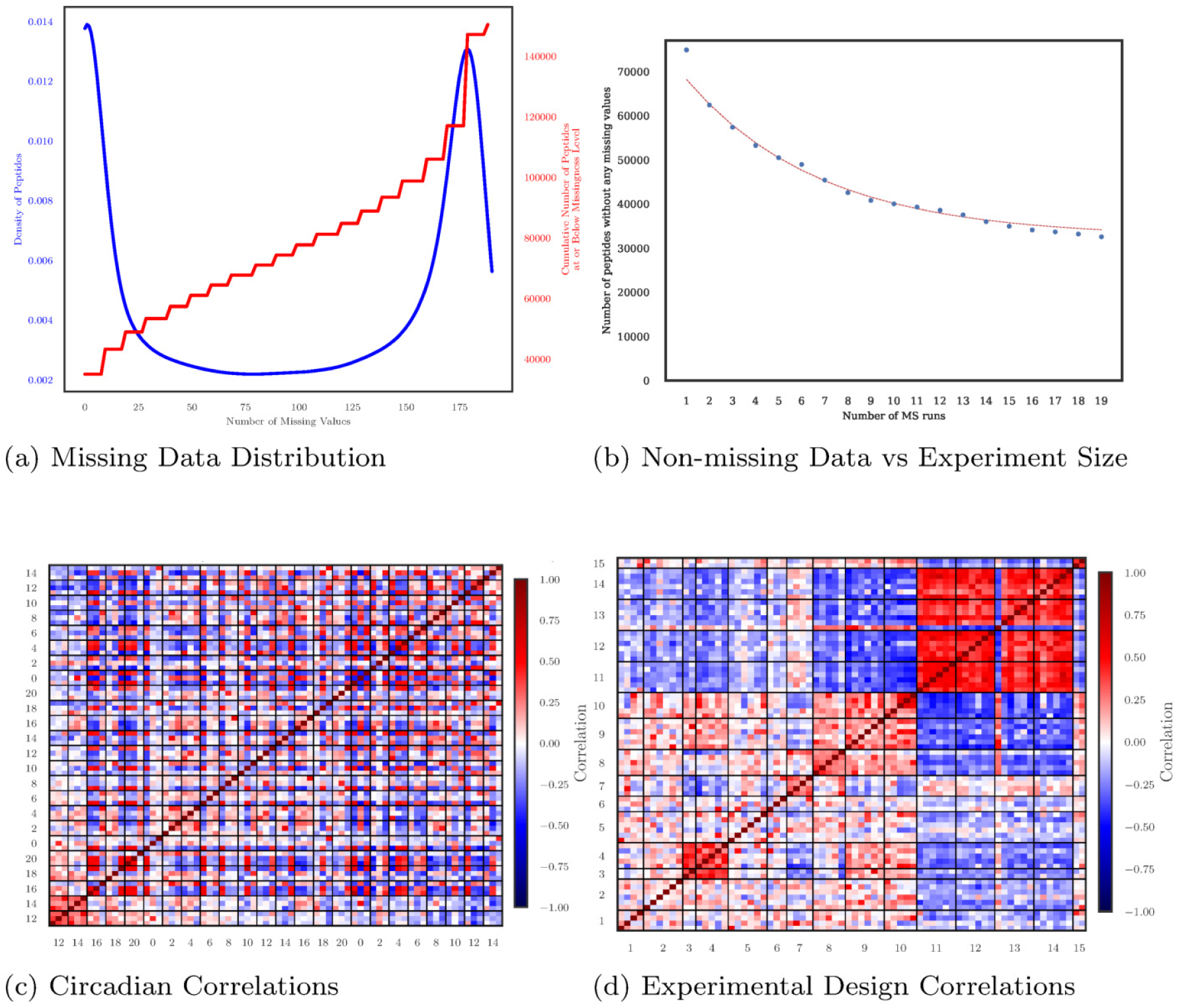
Illustration of dataset issues addressed by LIMBR (a) Distribution of missing data is shown in blue with the number of missing values on the x axis and the Kernel Density Estimate of peptides with that number of missing values on the left axis. The number of peptides observed with fewer than the given number of missing values on the x axis is shown in red with corresponding values on the right axis. (b) Number of peptides with no missing values (complete cases) as the number of MS runs is increased. For a complete case analysis, the number of peptides considered decreases exponentially as the size of the experiment is increased (exponential fit shown in red). (c) Correlation matrix of WT experimental data. The matrix is sorted by the time the sample was taken and the triplicate structure is indicated by the 3×3 black boxes. Numbers indicate the circadian time of the observation. Minimal Correlation is observed between replicates and between samples with matched circadian times. (d) The same correlation data as in C is here shown sorted by the sample preparation set. Strong correlations are seen within sets, indicating that sample preparation was a major contributor to the observed signal.

Bias trends that correspond to batch effects were also evident in this large experiment. We sorted samples by time point (Fig. 1 c) and did not observe evident patterns associated with time, which we would have expected because of the nature of circadian regulation in which expression exhibits gradual cyclical changes. For a dataset without batch effects, we would observe strong agreement between replicates and a larger checkerboard pattern, as day-phase protein expression should have been similar and generally opposed to night-phase expression. Instead, arranging samples by preparation set, which indicated the batches to which samples were randomized for purification, digestion and other processing prior to the M/S run, revealed pronounced patterns (Fig. 1 d). In a dataset without batch effects we would expect to see no discernible structure in this plot.

LIMBR was developed to model and remove bias trends in this type of data (Fig. 2). After missing data was imputed, batch effects were modeled using an SVA-based method. By modeling replicate and time series correlations, we produced a matrix of residuals containing the unknown batch effects. We modeled these batch effects with Singular Value Decomposition (SVD) to produce linearly independent bias trend. By permuting the residual matrix, we estimated the significance for each bias trends and removed those passing a user-specified significance threshold. The LIMBR software implements these methods alongside imputation methods to provide a toolkit for the analysis of large proteomic experiments.

**Fig. 2.**
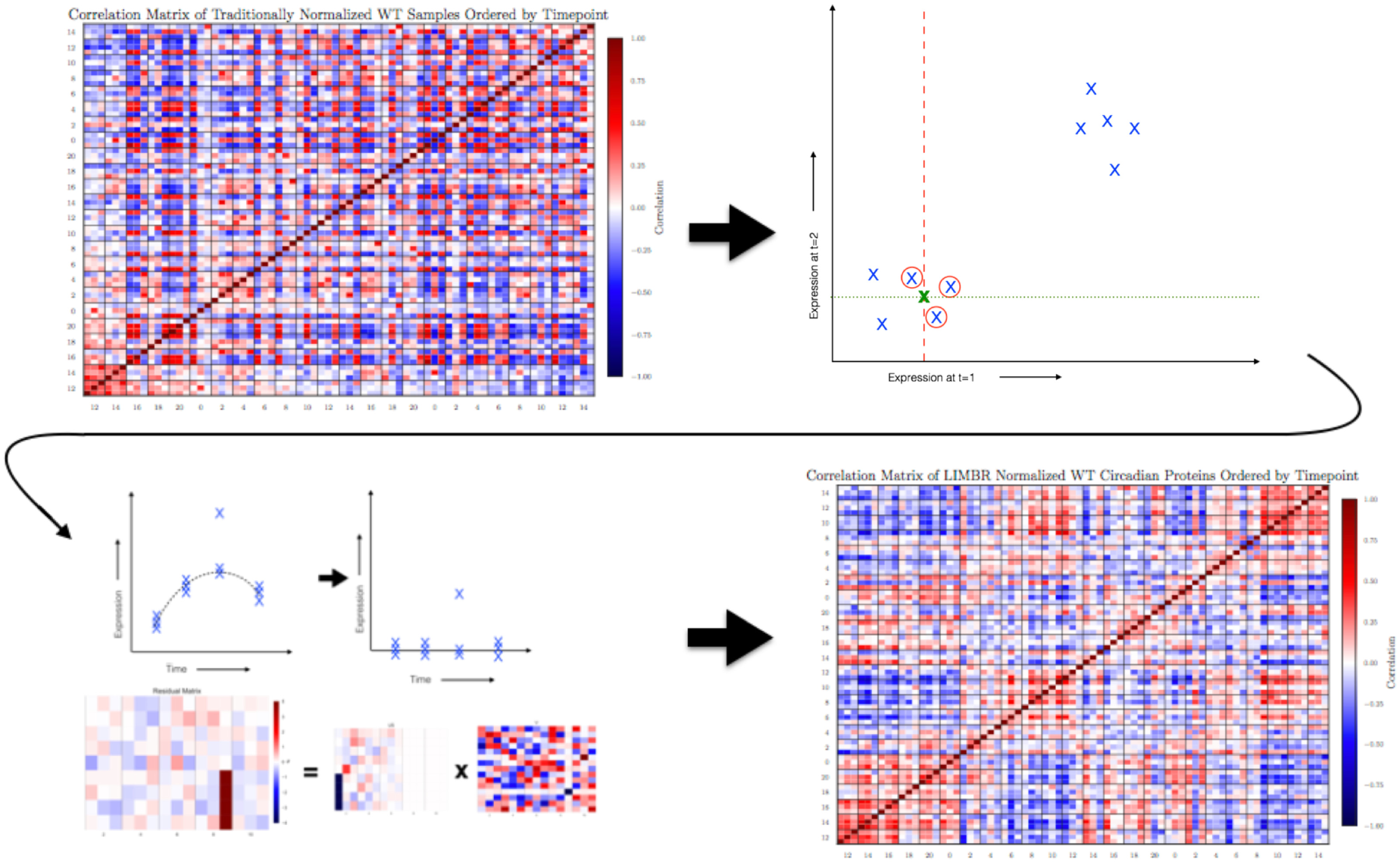
Algorithm Outline The outline of processes performed by the LIMBR algorithm is diagrammed, beginning with raw data heavily influenced by bias trends, proceeding to imputation of missing data, modeling of bias trends and the final removal of those bias trends producing a denoised dataset.

## Methods

### Imputation of Missing Values

In the proof-of-concept dataset only 32,561 peptides were measured in all samples; however, 137,340 peptides were measured in at least one sample. We sought to increase the coverage of samples to avoid the situation where increased numbers of replicates led to fewer analyzable peptides. For this, we implemented K nearest neighbors imputation (KNN), which has been found to be fast and reliable at the imputation levels and dataset sizes relevant to such proteomics experiments Batista and Monard, 2001; Troyanskaya *et al*., 2001; Wasito and Schmitt, 2005; Mandel *et al*., 2015.

The KNN algorithm imputes missing data by finding the K nearest data points with complete data for a given data point and imputes the missing value as the average of the nearby points’ values. Where *n* is the number of observed peptides, *m* is the number of MS experiments, *𝒟*^*n×m*^ is our dataset and *d* is a datum (all observations on a peptide), for *d ∈ D* we found the **K** nearest neighbors of d by Euclidean distance. We then constructed the matrix *𝒩* ^**K***×m*^ with rows being nearest neighbors of d. Finally we imputed the missing values in d as the mean of the corresponding columns of *𝒩* (second panel Fig 2). We report results using 10 nearest neighbors, which has been shown to be appropriate for datasets of roughly this size Wang *et al*., 2017b. In light of the triplicate experimental design employed for this data and the fact that KNN has been shown to function effectively on biological datasets without replicates up to 35% missingness, we imputed peptides missing in fewer than 30% of samples Mandel *et al*., 2015. Both parameters may be set at runtime in LIMBR by the user. An exploration of the impact of parameter selection on the analysis of simulated data is provided in figure S1. Values for k in the range of 5-15, along with missingness thresholds in the 30-40% range are recommended.

Even with this conservative imputation threshold, applying KNN increased the size of the proof-of-concept dataset by approximately 70% (32,561 peptides to 56,353 peptides), substantially improving both the number of proteins detected and eliminating the decrease in analyzable peptides with increasing numbers of replicates.

### Modelling of Bias Trends

MS experiments are subject to complex batch effects which can contribute a sizeable fraction of the total variability recorded in an experiment. Because circadian time courses require many time points and replicates and the biological trends are relatively subtle, these bias trends are often stronger than the biological signal in such datasets.

We first recognize that some peptides will be more closely correlated with the process of interest to investigators and that our modeling of bias trends can be improved by excluding such peptides Leek, 2007. In the case of circadian experiments we use the ratio of autocorrelations at 12 and 24 hours as a measure of correlation (*c*) to our variable of interest (circadian expression). The correlation threshold at which peptides are not analyzed for batch effects (*c*_*t*_), can be specified by the user and should be set relatively low (we use 25%) to screen out only those peptides where the signal of interest greatly outweighs any batch effects. In this way, we formed the reduced data matrix, *𝒟*_*r*_ of peptides for which *c < c*_*t*_.

Next, we proceeded to separating batch effects from our dataset. For a time series experiment, we expect some simple correlations to describe the data. Namely, we expect that replicates will agree with one another and that adjacent timepoints will be more similar than distant ones. Given a series of timepoints for each observation S, this structure is captured mathematically by a LOWESS model which we fitted to our data:

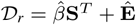

We then analyzed the residuals, **Ê** from this model calculated as:

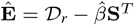

These residuals represent variability in the data set which did not agree with the experimental design and should therefore contain any batch effects. Since completely eliminating all of the model residuals would have drastically underestimated our uncertainty, we chose to model batch effects within the residual matrix Jaffe *et al*., 2015; Leek, 2007. We did this by breaking the residual matrix **Ê**, down into into a set of linearly independent bias trends with singular value decomposition **Ê** = **UDV**^*T*^. SVD has been used to model bias trends in micro-array, RNAseq and M/S data Parsana *et al*., 2017; Tsang *et al*., 2014; Karpievitch *et al*., 2009. Once we had broken down the residual matrix into bias trends, we needed to determine which were likely to have been contributed by batch effects and should therefore be removed from the dataset. We did this by estimating which of these trends contributes more variability than we would expect at random. By repeatedly permuting the residual matrix and performing SVD, we estimated the null distribution of variance explained by each singular vector and thereby assessed the ‘significance’ of individual bias trends, removing only those which explain more variance than we would expect by chance at a user specified threshold *p*. We designated bias trends with *p <.*05 as significant. With *d*_*l*_ as the *l*^*th*^ singular value, for right singular value k = 1, …, *n* we calculated the observed test statistic *T*_*k*_ as:

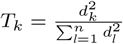

We then permuted each row of the matrix **Ê** independently to form a matrix **Ê**\*. We calculated the singular values and the null statistic 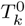 in similar fashion 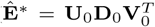 and 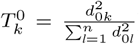). These calculations were repeated *B* times and the p value for right singular vector k was calculated as:

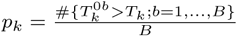

Having calculated the significance of each bias trend, we could have simply removed those passing a given threshold. Instead we chose to further remove our modeling choices from the bias trends we calculated Leek, 2007. We did this by moving through the trends in order adding one for each trend where *p*_*k*_ *< α* and stopping when this condition is no longer met, thereby estimating the number of significant trends *sb* as:

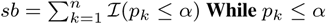

We then regressed each significant trend **v**_*k*_ against the rows of *𝒟*_*r*_ calculating a p value for their association. Given an estimate of the background uniform distribution of p values 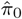, the number of truly associated peptides for each trend, 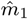, was estimated as:

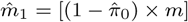

A subset of the peptides with the 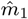 smallest p-values for that trend was then formed. The batch effect *j*\* was then modeled as the right singular vector of the reduced subset matrix most correlated with the bias trend being modeled:

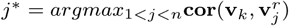

where the right singular vectors of the reduced subsetted matrix are:

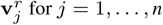

The matrix of batch effects **Ĝ**_*k*_ was then taken to be 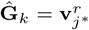 and these batch effects were removed by returning *𝒟* − **M** *×* **Ĝ** where **M** is found by a least squares solution of *𝒟* = **M** *×* **Ĝ**. By first reducing the data considered to only those peptides not correlated with our variable of interest, we avoid learning circadian expression patterns as a bias trend. We then quantify the variability contributed by the linearly independent bias trends produced by singular value decomposition of the residual matrix. By repeatedly randomizing the residual matrix and re-performing those calculations we estimate the likelihood of each bias trend contributing the variability observed by chance and can label some bias trends as significant based on this estimate. We then reduced the impact of the already limited modeling assumptions we had employed by regressing the significant trends against the original data and calculating our bias trends from a subset of that data most correlated with the original trends. In this way we were able to infer and remove batch effects without knowledge of the sample handling steps that produced them.

### Simulation Studies

To verify the effectiveness of the batch effect modeling and removal procedure, we conducted simulation studies. First, we generated synthetic data with 50% circadian peptides taken from sine waves equally distributed between opposite phases and added gaussian random noise. The other 50% of peptides were non-circadian and generated solely from gaussian random noise. We added randomly generated batch effects to these data and processed the resulting datasets with LIMBR. We analyzed the datasets before noise (Fig. 3 a panel 1), after the addition of noise (Fig 3 a panel 2) and after processing with LIMBR (Fig 3 a panel 3) with eJTK-cycle. We compared the resulting circadian classification accuracies using ROC curves.

**Fig. 3.**
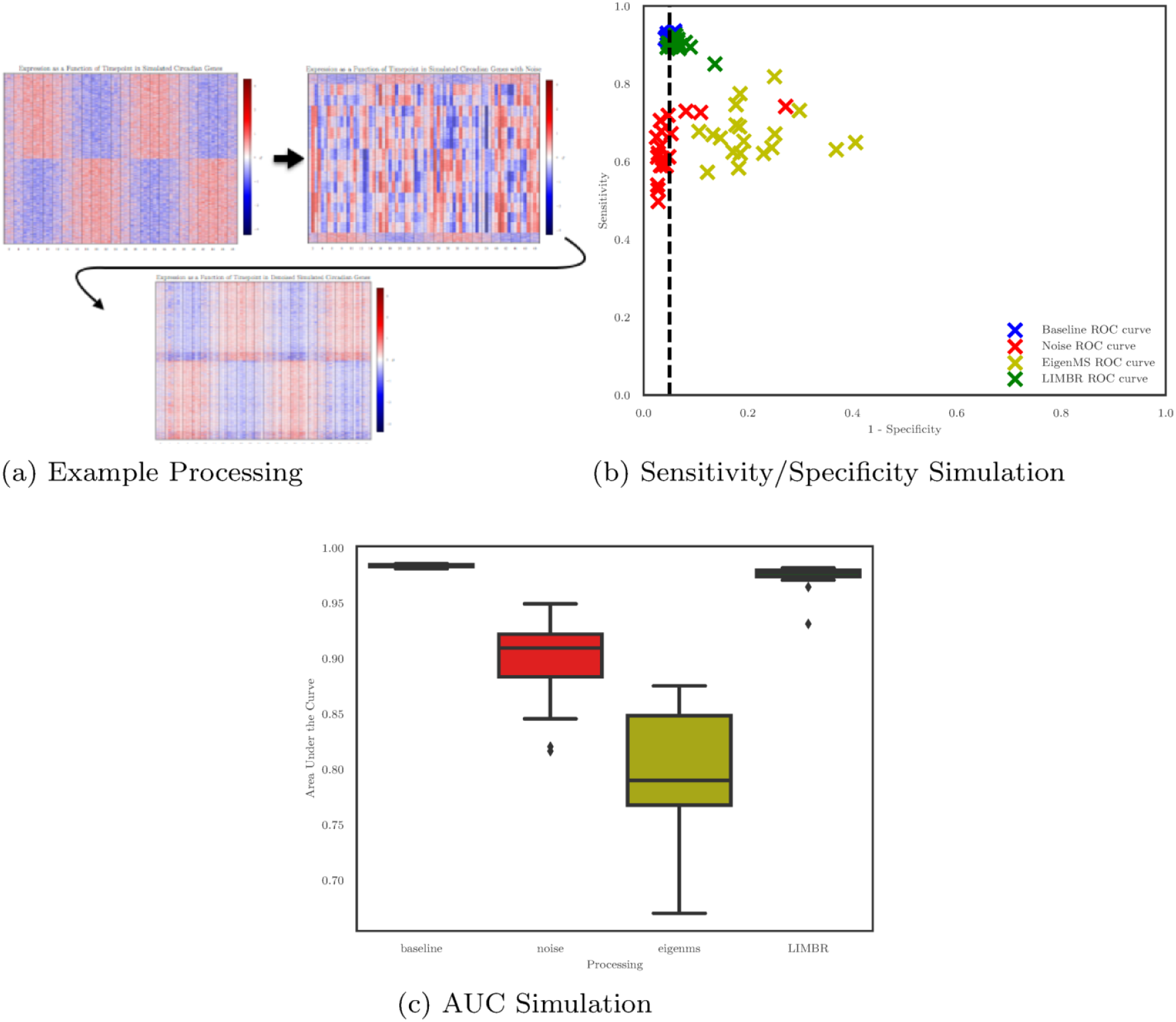
Simulation Results Example results showing initial circadian signal, data with the addition of 3 random bias trends and data after the application of LIMBR. (b) Representative sensitivity and specificity data showing the effectiveness of circadian classification for twenty rounds of simulation using a *p <.*05 threshold in eJTK for each of the three stages of data processing shown in (a) with the alternative eigenMS method shown for comparison. (c) Area under the receiver operating characteristic curves for the same twenty simulations shown in (b).

In order to provide a more stringent evaluation, we conducted an additional set of simulations in which three randomly generated batch effects were applied to the simulated data. Since the batch effects were applied at random, this resulted in many fewer unaffected peptides (*∼* 12.5% vs *∼* 50%) and many more combinations of bias trends (7 vs 1) making this data much more challenging to analyze. Both the single batch effect and triple batch effect simulations were repeated a total of twenty times and the results for the more challenging simulations are visualized in Fig 3 b and 3 c. As an additional point of comparison, we processed the triple batch effect simulations with eigenMS the current state of the art sva based software for removing batch effects in MS data Karpievitch *et al*., 2009. The addition of three bias trends completely obliterated circadian classification accuracy. Processing with eigenMS, which cannot exploit the time series structure of the dataset was largely ineffective. Processing with LIMBR; however, almost completely recovered classification accuracy. These simulations indicated that LIMBR was modeling and removing unknown bias trends accurately and reproducibly.

### Application to Biological Data

Since the two genotypes were randomized and run on the M/S together, the batch effects observed in each dataset should be the same. Additionally, while we do expect some differences in expression between the two, globally expression should be similar. When analyzing the data however, we applied LIMBR to the time courses from each of the two genotypes independently. Since information was not shared with the algorithm between these runs, comparing the results of these analyses allows us to examine the reproducibility of our techniques.

First, we compared the bias trends identified by LIMBR with the sample handling steps performed prior to M/S. We found that for both genotype time courses, the bias trends correlated well with sample preparation steps (Fig. 4 a and 4 b). The primary sources of bias in sample preparation were strongly correlated with sample preparation and TMT sets indicating that these steps made major contributions to the observed bias. This fits well with previous reports that peptide digestion and charge state are major sources of bias in M/S quantification Rudnick *et al*., 2014; Piehowski *et al*., 2013. We then compared the bias trends found in each genotype, finding that LIMBR had independently reproduced analogous bias trends for each dataset (Fig. 4 c).

**Fig. 4.**
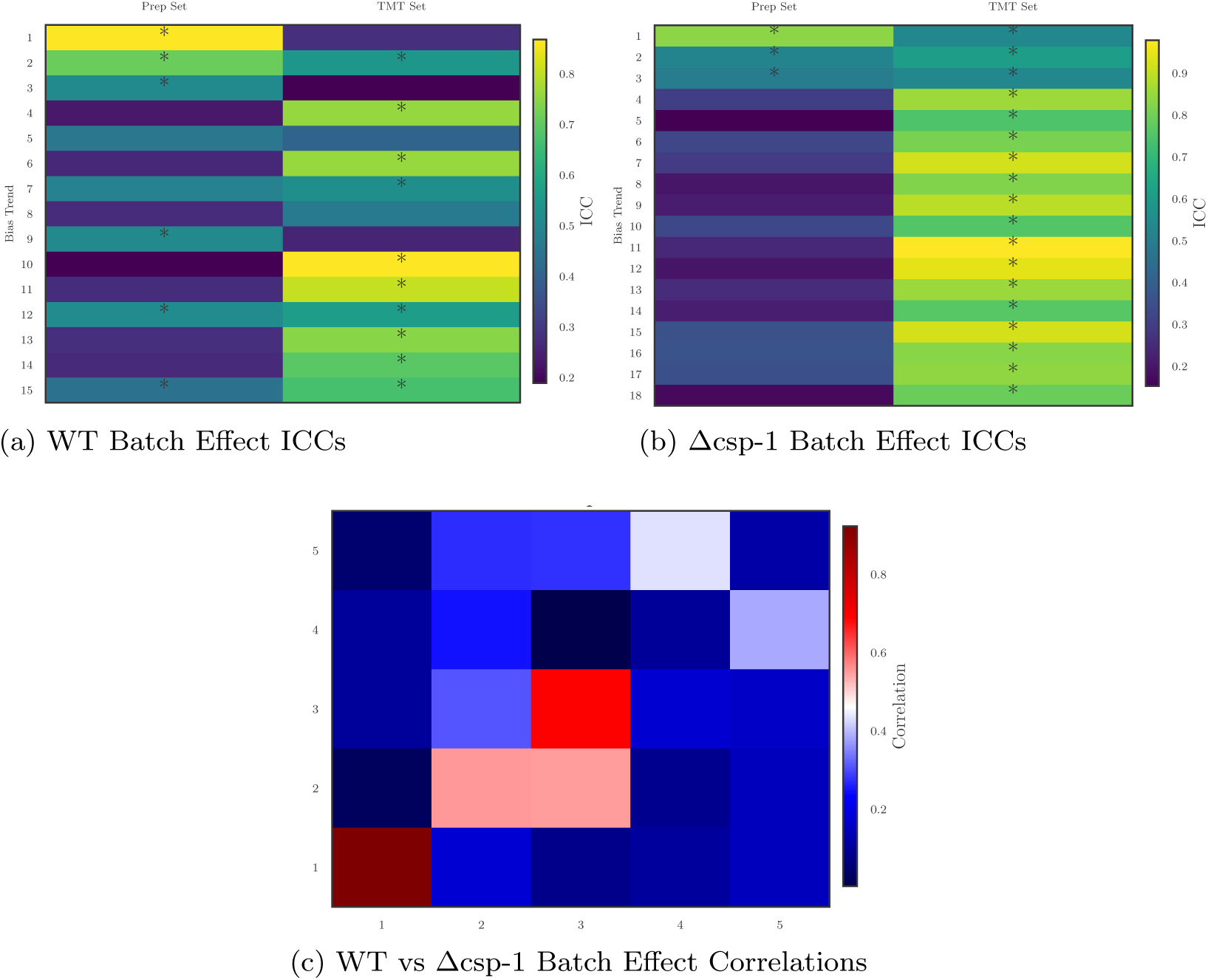
Analysis of Batch Effects Identified in Biological Data Interclass correlation coefficients between batch effects identified as significant and the prep set and TMT set of each sample in WT (A) and Δcsp-1 (B) are shown with * indicating significance at the *p <.*001 level for a one way F-test. Almost all bias trends are significantly associated with sample preparation or TMT labelling sets. In some cases, due to incomplete randomization, bias trends are associated with both. C. Correlation matrix of the top 5 bias trends as measured by explained variance in the WT and delta csp-1 experiments. The top 3 bias trends independently identified in each experiment are quite similar, and trends 4 and 5 seem to be flipped in their rankings but still correspond closely.

After these final evaluations of the software, we analyzed the results of circadian classification for our LIMBR processed data and found 324 circadian proteins out of 4754 analyzed proteins (7%) with a predominantly bimodal distribution of phases (Fig. 5 a). Analysis of the sample correlation matrix for the WT dataset revealed a much stronger agreement between replicates along with the diurnal checkerboard pattern (Fig. 5 b). When we focused the analysis on only circadian proteins, these patterns were even more pronounced (Fig. 5 c).

**Fig. 5.**
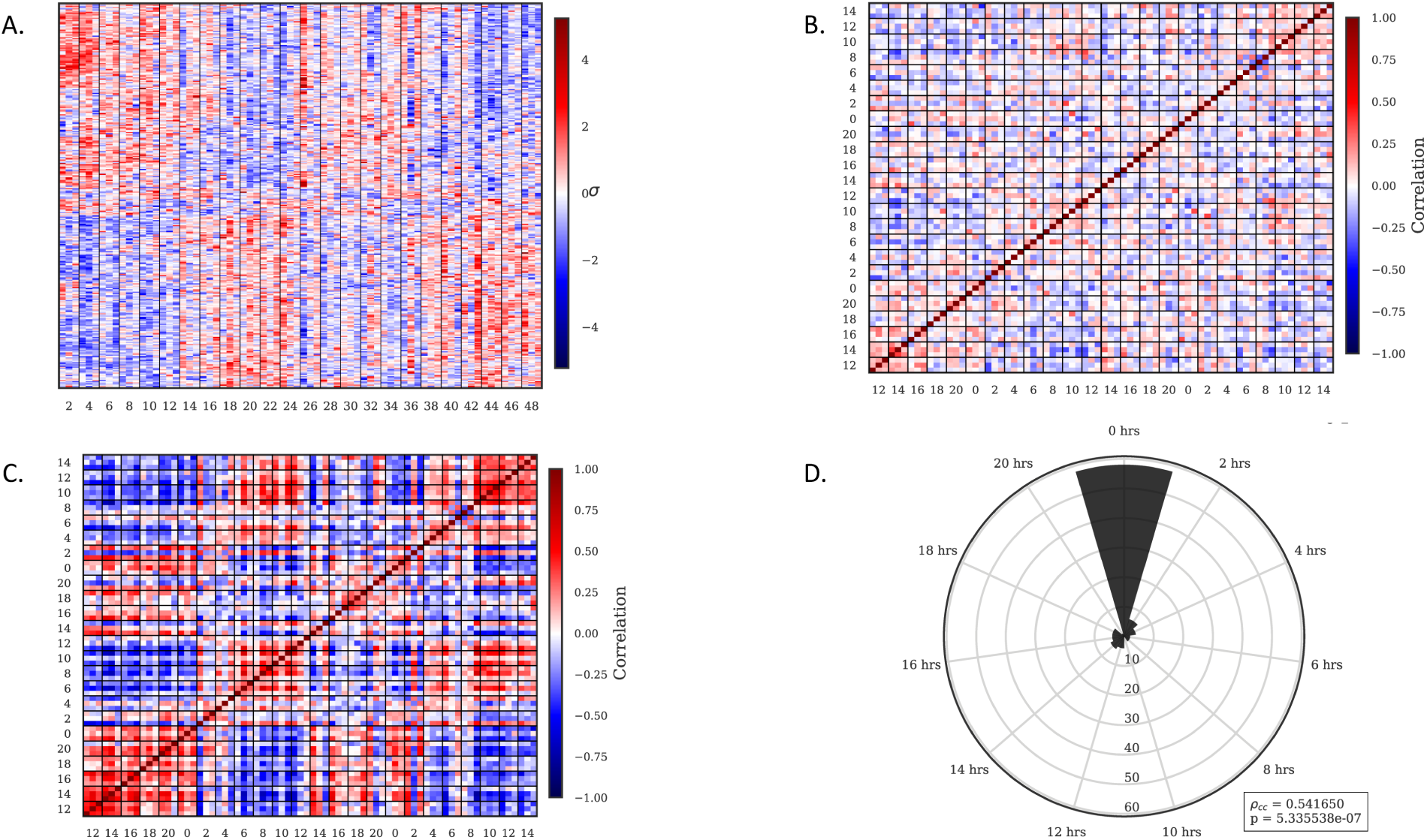
Results of application of LIMBR to biological data A. Expression profiles of genes identified as circadian in the WT dataset after processing. B. Correlation matrix of WT dataset after processing, showing increased similarity between replicates and checker boarding pattern of similarity between repeated circadian times (compare to figure 1B). C. Correlation matrix of WT dataset generated only from proteins identified as circadian showing pronounced checker boarding similarity between repeated circadian times. D. For proteins identified as circadian in both genotypes the distribution of differences in predicted phases between the genotypes is shown. A strong peak at 0 indicates that the predicted phases of these genes match well between the independent processing runs.

We compared the predicted phases of proteins identified as circadian in both genotypes and found a tight correspondence between the two (Fig. 5 d). A detailed discussion of the biological implications of these analyses is underway (Hurley et al., in preparation).

## Discussion

The combination of KNN based imputation and SVD based error modeling improve the results of time-course proteomics analysis markedly. These techniques, which we implement into the LIMBR software package, allow for the recovery of circadian signals in simulated and biological datasets for which no such signal is initially observable.

Simulation studies reveal that LIMBR has high sensitivity and specificity along with effective recovery of phase. LIMBR also produces superior results to other sva based methods for the removal of batch effects when applied to time series data. This increase in performance can be attributed to the reduction and subsetting steps, but especially to the LOWESS based calculation of residuals. The reduction step serves to prevent learning signal of interest as a batch effect, while the subsetting step helps to draw batch effects more directly from the data with less influence from our modeling decisions. The LOWESS based calculation of residuals allows for an improved estimate of batch effects by incorporating experimental structure which is lost when processing with eigenMS. While the performance improvements of LIMBR over eigenMS for the simulated data in this study are drastic, this difference can be attributed to the much greater amount of information LIMBR has access to for time series data. LIMBR should therefore be expected to offer larger improvements over other methods for time series data than for block experimental designs where LIMBR’s improvements in residual calculations do not come into play. Even in the case of a smaller block design experiment with relatively fewer and smaller batch effects however, LIMBR still offers the added benefit of integrated imputation of missing values.

In the case of biological data, we also show that LIMBR produces effective and consistent results. When comparing between proteomics datasets for two genotypes, we see that the inferred batch effects match well with experimental parameters. This indicates that the batch effects being removed are well grounded in reality. Additionally, batch effects inferred from independent processing of the two genotypes match well with one another, indicating the consistency of LIMBR. Finally, the phases of circadian proteins found in these independently processed datasets closely agree. The consistency of batch effects with experimental parameters and between LIMBR runs along with the agreement of phases of circadian genes between LIMBR runs indicates the consistency of our methods, while the recovery of circadian signal from a dataset otherwise dominated by batch effects speaks to the efficacy of these techniques.

While it is possible for the removal of bias trends such as batch effects to inflate false positive rates in studies of differential expression, these effects should only be a concern when there is a highly uneven distribution of samples to batches Nygaard *et al*., 2015. Additionally the application of bias modelling techniques has been shown to increase the sensitivity and specificity of differential expression studies by removing bias trends and to be a net positive when applied to RNAseq data Li *et al*., 2014. The algorithm employed here is based on earlier work but is notable both for being a modern implementation in python and for implementing a more advanced reduced subset method originally proposed by Leek in tandem with LOWESS based calculation of residuals Leek and Storey, 2007b; Leek *et al*., 2012.

## Conclusion

The application of LIMBR to this dataset demonstrates the potential these techniques hold for the analysis of large scale circadian proteomics time courses. In three separate prior studies of circadian proteomics, Mauvoisin *et al*., 2014; Robles *et al*., 2014, 2016 the issues of lost data resulted in circadian data for less than 200 proteins, even after accepting time courses in which many observations were missing and where rhythmicity was called with an FDR of 0.25 to 0.3. With LIMBR, we can apply a much lower FDR of.05, reject data sets in which more than 30% of the points are missing, and still identify many more circadianly regulated proteins. While this highlights the strengths of the technique, it is also worth noting the limitations, specifically that randomization of samples during MS as well as a high number of time points and replicates are critical to effectively model batch effects. Additionally, randomization of samples to MS runs and processing steps must be performed independently and the standard practice of pool normalization should be employed to ensure that batch effects do not coincide with missing data, thereby biasing the imputation step. While such large experiments require the investment of additional resources, their importance to this type of study cannot be overemphasized. Although there is still room for improvement in the design of circadian proteomics experiments both in vitro and in silico, this work makes clear the need for analysis techniques and software for such datasets.

## Availability

Python code and documentation is available for download at https://github.com/aleccrowell/LIMBR and LIMBR can be downloaded and installed with dependencies using ‘pip install limbr’

## Algorithm 1 K Nearest Neighbors Imputation

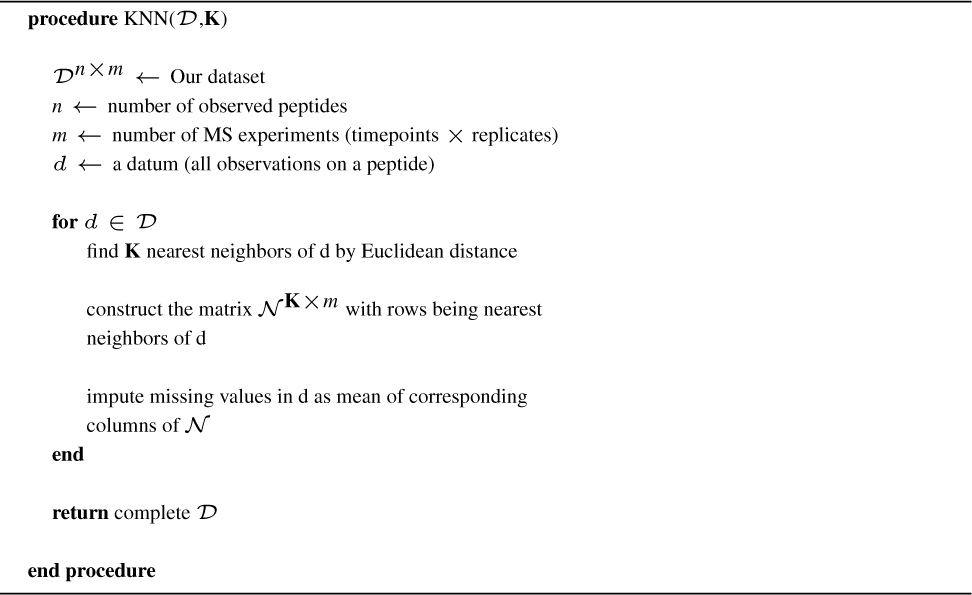

## Algorithm 2 SVD Bias Modelling

**procedure** SVD_Bias(*D, α*)

*𝒟*^*n×m*^ *←* Our dataset

*n ←* number of observed peptides

*m ←* number of MS experiments (timepoints *×* replicates)

*α ←* bias trend significance threshold

**S** *←* Sample timepoints

*c*_*t*_ *←* correlation threshold

calculate the correlation to the primary variable of interest for each peptide *c*

form a reduced data matrix *𝒟*_*r*_ of peptides for which

*c < c*_*t*_

fit the Lowess model *𝒟*_*r*_ = *β***S**^*T*^ + **E**

Calculate the residual matrix as **Ê** = *𝒟*_*r*_ *− β***S**^*T*^

Calculate the singular value decomposition of the residual matrix **Ê** = **UDV**^*T*^

With *d*_*l*_ as the *l*^*th*^ singular value, for right singular value k = 1, …, *n* calculate the observed test statistic as:

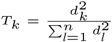

Permute each row of the matrix **Ê** independently to form a matrix **Ê**\*

Calculate the singular value decomposition of the matrix 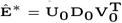

For right singular value k calculate the null statistic:

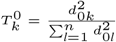

Repeat calculation of the null statistic B times

Calculate the p value for right singular vector k as:

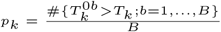

Estimate number of significant trends *sb* as:

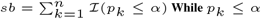

For each significant trend **v**_*k*_, regress **v**_*k*_ on each row of *Dr* calculating a p-value for their association

Estimate the number of truly associated features as 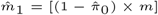 and form a subset of features with the 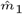 smallest p-values

Calculate the right singular vectors of the reduced subsetted matrix as 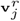 for *j* = 1, *…, n*

Estimate the surrogate variable *j*\*as the eigengene of the reduced subset matrix most correlated with the corresponding residual eigengene

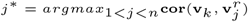

Set 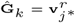

Find least squares solution for **M** where *𝒟* = **M** *×* Ĝ

**return** *𝒟* − **M** *×* Ĝ

**end procedure**

